# EEG connectivity changes in early response to antidepressant treatment

**DOI:** 10.64898/2026.03.18.712812

**Authors:** Aditi Kathpalia, Ioannis Vlachos, Jaroslav Hlinka, Martin Brunovský, Martin Bareš, Milan Paluš

## Abstract

**Objective:** Finding indicators of early response to antidepressant treatment in EEG signals recorded from patients suffering from major depressive disorder.

**Methods:** Functional brain connectivity networks based on weighted imaginary coherence and weighted imaginary mean phase coherence were computed for 176 patients for 6 different EEG frequency bands. Cross-hemispheric connectivity (*CH*) and lateral asymmetry (*LA*) were estimated from these networks based on EEG signals recorded before the beginning of treatment (*V is*1) and one week after the start of the treatment (*V is*2). Repeated measures ANOVA was used to check for statistically significant changes in connectivity based on these measures at *V is*2 w.r.t. *V is*1. Post-hoc analysis was performed with multiple pairwise comparison tests to determine which group means were significantly different.

**Results:** It was found that *CH*_*V is*2_ was significantly reduced w.r.t. *CH*_*V is*1_ in the *β*_1_ [12.5 - 17.5 Hz] frequency band for the responders to treatment. Also, *LA*_*V is*2_ was significantly increased w.r.t. *LA*_*V is*1_ in the *β*_1_ frequency band for the responders. No such significant changes were observed for the non-responders. Brain networks constructed using both weighted imaginary coherence and weighted imaginary mean phase coherence were found to exhibit these results. For the CH connectivity changes, binarized networks and for the LA connectivity changes, weighted networks were found to be more reliable.

**Conclusions:** Responders were found to show a reduction in cross-hemispheric connectivity and an increase in lateral asymmetry, both in the *β*_1_ band while no such change was observed for the non-responders.

**Significance:** Decrease in cross-hemispheric connectivity and increase in lateral asymmetry in the *β*_1_ band may represent candidate neurophysiological indicators of early treatment response, but they require independent replication before any clinical application can be considered.

## 1. Introduction

Major Depressive Disorder (MDD) is one of the most common psychiatric illnesses of the world. WHO ranked it as the third highest cause of the burden of disease worldwide in 2008 and has expected it to become the first highest cause by 2030 (WHO, 2008). MDD can often be chronic and its patients don’t easily respond to treatment, often undergoing several rounds of trial and error for finding effective treatment. Hence, to be able to predict the response to a given treatment early on can be helpful in disease management and control. Particularly, it can help to save time, resources and also reduces the risk of deteriorating patients’ clinical status and their subsequent loss of interest in seeking and complying with the medication or therapy being administered.

While various neuroimaging, neurophysiological and other markers have been studied in relation to and found to be correlated with antidepressant response (Gudayol-Ferré et al., 2015; Stade and Iosifescu, 2016), none of them have reached the status of being used in the clinical practice for prediction purposes (Breitenstein et al., 2014). The study of electroencephalography (EEG) based biomarkers is the most common in this domain naturally due to its relative ease of acquisition, non-invasiveness, relatively low cost, wide availability and consequent wide applicability (Iosifescu and Lapidus, 2011). However, neither of these as well, is close to being used as a clinically proven predictor.

From the past couple of decades, analysis of individual EEG electrode signals and their characteristics such as spectral power and complexity have served as potential biomarkers for several disorders (Neto et al., 2015; Riaz et al., 2015; Bosl et al., 2011) including depression and its treatment (Hinrikus et al., 2010; Li et al., 2008; Bares et al., 2015b,a, 2019). In the recent years, brain connectivity analysis has become an important tool to understand different brain states and disorders (Lang et al., 2012; Friston, 2011). Brain connectivity can be studied at the level of anatomical connections and this is referred to as structural connectivity. It can also be studied based on construction of functional brain networks by estimating the functional dependence/connectivity between signals recorded from different brain regions, and this is referred to as functional brain connectivity. In case of EEG, this is most commonly done pairwise for signals acquired from the electrodes placed on scalp. In several studies, it has been found that connectivity strengths obtained from different brain regions could be useful indicators of depression (Anand et al., 2005; Fingelkurts et al., 2007; Lee et al., 2011). Since normal brain function involves a lot of information transfer between its different regions, a dysfunction may be related to ‘abnormal’ information exchange between these regions and need not be localized to specific regions.

In this work, we analyse EEG recordings taken from patients undergoing depression treatment to **identify neurophysiological indicators for early response to antidepressant treatment**. Based on resting state EEG recordings, functional brain networks are constructed for each patient before the start of the treatment (Visit 1) and one week after the treatment began (Visit 2). Measures based on brain connectivity are computed for the two visits and statistical tests are performed to check if there are any significant differences in the estimates at Visit 2 w.r.t. Visit 1, while controlling for factors such as the type of treatment administered - pharmacological vs neurostimulation; as well as the type of response of the subject - *responder* vs *non-responder* (as determined after completion of the treatment). If significant changes in the computed measures arose as a result of early response to anti-depressant treatment, we would expect to see changes in the group means (of the values for visits) for the responders alone and not the non-responders.

*Coherence* is conventionally used to study EEG-based brain connectivity in different spectral bands. However, since most experiments record scalp EEG with a common reference electrode as well as due to volume conduction effects, the presence of common signal can be found at all EEG electrodes. This can lead to spurious connectivity being observed in any given pair of channels (in any spectral band). Hence, in this work we use *weighted imaginary coherences* as well as *weighted imaginary mean phase coherences* to estimate connectivity. These measures are known to control for conductiv1ity effects and present ‘true’ physiological connectivity^1^ in any given pair of channels in the different EEG frequency bands considered (Paluš et al., 2023; Nolte et al., 2004; Pascual-Marqui et al., 2011). The estimation of the chosen measures was based on wavelet decomposition, instead of fourier decomposition, as wavelets are known to provide reliable estimates in case of EEG signals which are nonstationary (Sifuzzaman et al., 2009; Sleziak et al., 2015). Based on the functional brain networks obtained, the measures (or features) that are computed for comparison are *cross-hemispheric connectivity* and *lateral asymmetry* in different frequency bands. Previous literature suggests that alterations in neural synchronization and network reorganization as a result of depression can manifest through changes in these features (Quraan et al., 2014; Hermesdorf et al., 2016).

This paper is organized as follows. Section 2 describes the EEG data used in the study, the functional connectivity measures used for computation of brain networks and the features extracted from these brain networks. Section 3 contains the statistical data analysis results answering the question of whether there is a significant change in the features at Visit 2 compared to Visit 1 and what are the factors that lead to this change. We conclude in Section 4 with a discussion and conclusions of the study.

## 2. Materials and Methods

### 2.1. EEG data

EEG data used in this study was obtained from the database of the Prague Psychiatric Center/National Institute of Mental Health, Czech Republic. It was a part of research projects that were reviewed and approved by the ethical committee of the institute. All procedures and study design were in accordance with the latest version of the Declaration of Helsinki. EEG recordings from 188 patients with MDD were included in the study. Complete details of the sample and participant recruitment criteria are available in Bares et al. (2010, 2015a,b).

EEG data were recorded using a 19-channel system based on the international 10–20 electrode placement standard, with an initial sampling rate of 250 Hz or 1000 Hz (the latter was downsampled to 250 Hz for consistency). Data acquisition was carried out using a standard 21 channel digital EEG amplifier BrainScope (Unimedis, Prague) equipped with Ag/AgCl electrodes. The amplifier digitized the signal with a 16-bit resolution and a sensitivity of 7.63 nV/bit ( 130 bits/mV), providing a dynamic range of 250 mV. Electrode impedances were maintained below 5 kΩ throughout the recordings. The 21 surface electrode recordings were referenced to the electrode located between the Fz and Cz electrodes in the midline (FCz).

Approximately 10 minutes of recording were obtained from each of the patients lying in a semirecumbent position with eyes closed in a maximally alert state. The room in which the recordings were taken was dimly lit and equipped with sound attenuation facility. Further, these EEG recordings were done in the morning (10:00-12:00am) to minimize circadian variability. Caffeine intake and other potential confounding substances were controlled prior to EEG recording. Participants were instructed to abstain from stimulants for a minimum of 12 hours before the session. Alertness was monitored throughout the recording. If signs of drowsiness appeared in the EEG, the EEG technician used acoustic stimuli to awaken the subjects.

Four weeks of antidepressant treatment was administered to the patients based on the decision of the psychiatrist. The treatment was done by either serotonin-norepinephrine reuptake inhibitors, transcranial direct current stimulation, repetitive transcranial magnetic stimulation, selective serotonin reuptake inhibitors, norepinephrine-dopamine reuptake inhibitors or another means. EEG recordings obtained were reviewed by experienced EEG technicians following established guidelines. 12 patients were removed from the analysis because either their recordings were distorted and not readable or the recordings for three or more channels were silent. This resulted in a dataset consisting of 176 patients in the age range of 18 to 65 years. The sex, age and response (to antidepressent treatment) distributions of these patients are listed in Table 1. Two EEG recordings, one before starting medication (referred to as Visit 1) and the other, one week after starting medication (referred to as Visit 2) were acquired from the patients. Strict electrode placement guidelines using the 10-20 system were followed, ensuring consistent electrode positioning across sessions.

**Table 1.**
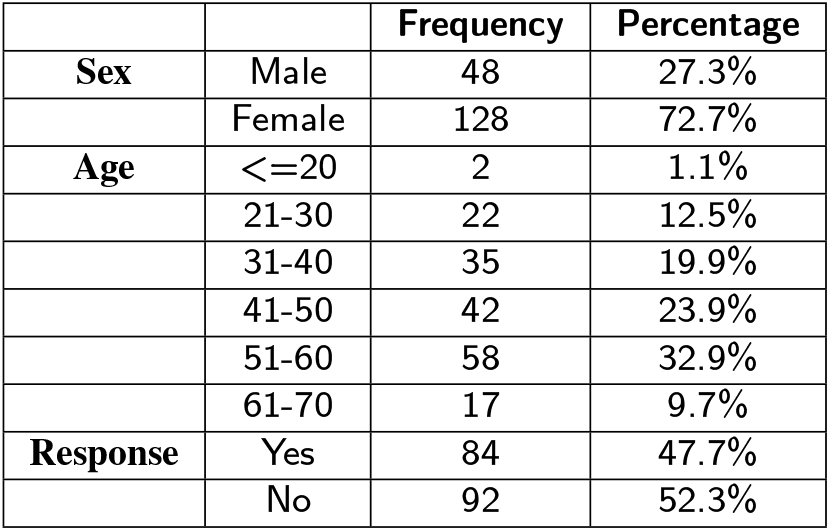
Demographic information of patients.

Montgomery and Asberg Depression Rating Scale (MADRS) (Montgomery and Åsberg, 1979; Williams and Kobak, 2008), a standardized, reliable, and valid questionnaire-based instrument, was employed by psychiatrists to assess the state of depressive symptoms in patients before start of the treatment and after the treatment period, which lasted for 4 weeks. The patients which thus responded/ not responded to treatment were decided on the basis of this MADRS score. For a patient to be considered a responder, the MADRS score should have reduced by at least 50% of the original score. Prior to participation, patients were fully informed about the study’s methodology and provided their informed consent for involvement.

### 2.2. Preprocessing

Signals from 19 standard EEG electrode positions were analysed for the study. These included Fp1, Fp2, F3, F4, C3, C4, P3, P4, O1, O2, F7, F8, T3, T4, T5, T6, Fz, Cz, Pz. Before the analysis, preprocessing of the signals was done. The EEGLab MATLAB toolbox (Delorme and Makeig, 2004) was used for data preprocessing. Since some recordings were made at 1000 Hz and others at 250 Hz sampling frequency, the recordings at 1000 Hz were subsampled to 250 Hz by taking every fourth sample. The first and the last 30s of the data were removed from the recordings as they may contain relatively more artifacts. Further, the EEG was transformed to average reference. This transformation minimizes reference-dependent biases in connectivity analysis. Consequently, the signals were band pass filtered from 1-40 Hz

Finally, the ‘clean windows’ function in EEGLab was used to remove segments of data with artifacts. This function cuts segments from the data which contain high-power artifacts. Specifically, only those windows of each patient’s EEG data are retained which have less than a certain fraction of “bad” channels. A channel is considered “bad” in a window if its power is above or below a given upper/lower threshold (set in terms of standard deviations from a robust estimate of the EEG power distribution in the channel). Parameters of the function were set such that the maximum number of bad channels allowed is 20% of the total number of channels present in the data. The window length used for assessing artifacts was set to 2s. This ensured that 2s segments of data were continuous when parsed from the beginning, despite the removal of artifacts. Raw and preprocessed EEG waveforms from channel Fp1 (that records activity from the prefrontal cortex) is depicted for a subject in figure 1 as an example.

**Figure 1:**
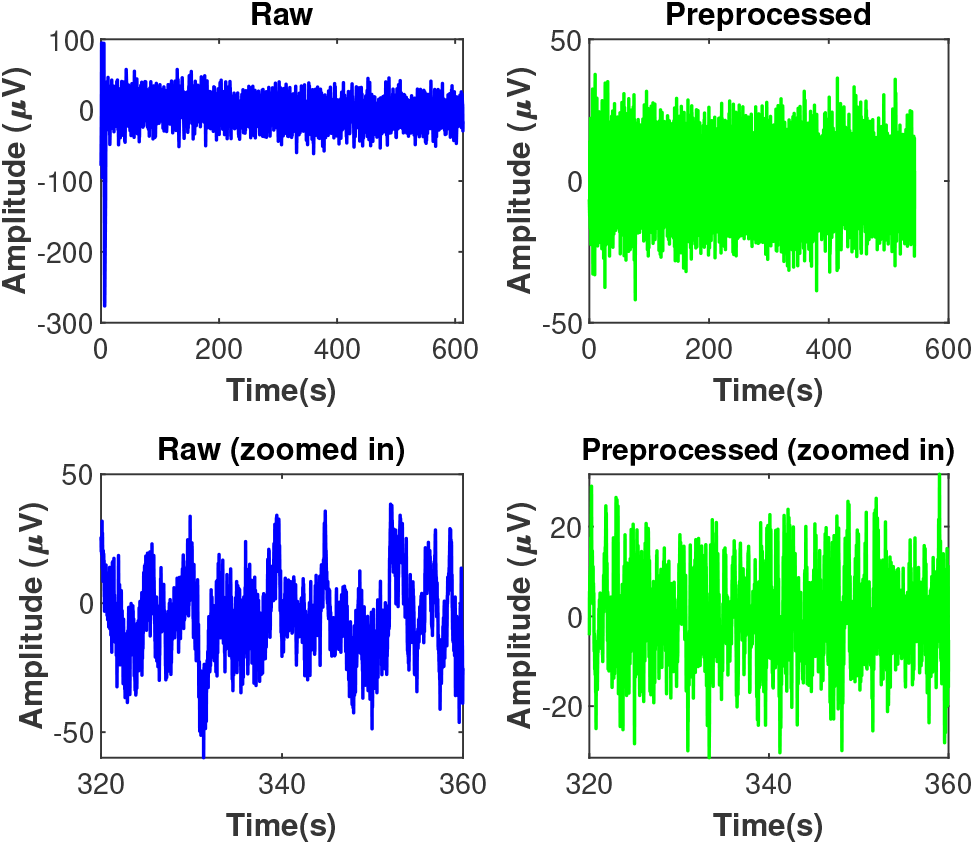
Raw and preprocessed EEG signals recorded from Fp1 electrode of a subject. Zoomed-in counterparts in the bottom row are plotted for better visualization.

### 2.3. Functional connectivity based analysis

For each patient, functional connectivity was quantified by estimating pairwise coherences from the preprocessed signals, for all EEG channel pairs. Brain networks constructed based on coherences for each patient were further used to assess connectivity changes that may happen in the responders soon after the start of the treatment. These were compared with the connectivity changes in the nonresponders.

#### 2.3.1. Wavelet-based Coherence estimation

The EEG signals from each of the 19 channels were first transformed into the wavelet domain using complex Morlet wavelet, yielding a set of complex wavelet coefficients. If *X*(*t*) is the signal from a channel, its wavelet coefficients are given by:

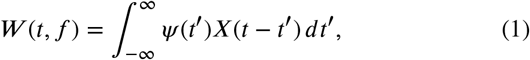

where, *ψ*(*t*) is the complex Morlet wavelet, given by:

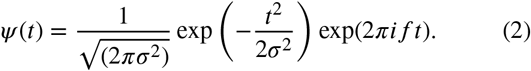

Morlet wavelet is essentially product of a complex sine wave and a Gaussian window. Here, *σ* is the width of the Gaussian, which depends on the time-frequency precision trade-off. Wavelet coefficients were computed for 78 different scales or frequencies, *f*, covering uniformly the range of 1-40 Hz. Using the complex wavelet coefficients *W*_*X*_(*t, f* ) and *W*_*Y*_ (*t, f* ) from signals *X*(*t*) and *Y* (*t*) recorded from two different channels, cross-wavelet coefficients can be estimated as:

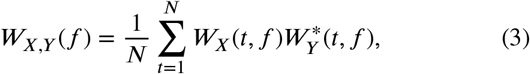

where 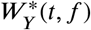 refers to the complex conjugate of *W*_*Y*_ (*t, f* ).

For the estimation of wavelet coefficients, we use the algorithm of Torrence and Compo (1998) (code available at http://atoc.colorado.edu/research/wavelets/) in which the optimized parameters are set as default. Since this setting leads to relatively narrow band wavelet components, instead of widening the bandwidth we computed wavelet based coherence for a number of narrow bands, results from which were integrated into conventional EEG frequency bands.

##### Weighted Imaginary Coherence (WIC)

Traditional coherence measures suffer from volume conduction effects as has been extensively studied and established in existing literature (Nolte et al., 2004; Stam et al., 2007; Srinivasan et al., 2007; Vinck et al., 2011; Sanchez Bornot et al., 2018). Hence, lagged connectivity approaches that rely on the imaginary part of coherence as in Nolte et al. (2004) and phase differences Stam et al. (2007) have been proposed.

In order to remove conductivity effects present between EEG signals while estimating pairwise connectivity, we adopted the formulation of ‘lagged coherence’ given by Pascual-Marqui et al. (2011). In Nolte et al. (2004), use of imaginary part of coherence was proposed as an index to get rid of instantaneous connectivity effects that may arise due to conduction. Pascual-Marqui et al. (2011) further suggested ‘weighing’ it by the instantaneous component of coherence such that their lagged coherence is less affected by low spatial resolution and contains almost pure physiological information. Hence, we call this index as Weighted Imaginary Coherence (WIC). This new nomenclature also helps to distinguish the index from another use of lagged coherence where coherence between a signal and its lagged version is estimated (Fransen et al., 2015).

Pascual-Marqui et al. (2011) compare their approach with other lagged connectivity approaches (such as in Nolte et al. (2004)) and show that it is able to robustly infer the presence of a lagged connection, particularly in the presence of high instantaneous connectivity. Additionally, for the EEG dataset used in this paper, the authors in Paluš et al. (2023) demonstrate how conductivity masks ‘true’ connectivity over all frequency bands and weighted imaginary coherence uncovers connectivity on physiologically relevant frequencies which is devoid of confounding effects.

Instead of Fourier coefficients used previously in (Pascual-Marqui et al., 2011) and subsequent papers, we use wavelets in the formulation of WIC. Wavelet based frequency decomposition shows better performance than Fourier for non-stationary and nonlinear signals (Sifuzzaman et al., 2009; Sleziak et al., 2015). Hence, wavelets are recommended for the analysis of EEG signals, which typically have these characteristics. Use of wavelets also enabled the estimation of time resolved brain connectivity networks.

Complex wavelet coherence can then be estimated as per Eq. 4 below:

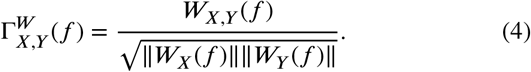

The WIC between channels *X* and *Y* at frequency *f* is estimated as:

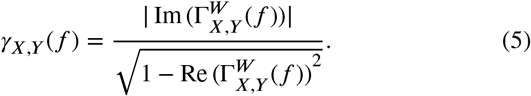

##### Weighted Imaginary Mean Phase Coherence (WIMPC)

The concept of phase synchronization between coupled chaotic systems was introduced in Rosenblum et al. (1996) and refers to an entrainment of the phases of the systems. It has been used in the analysis of physiological signals and also EEG signals in particular. One of the measures by which it is quantified is Mean Phase Coherence (Mormann et al., 2000; Mezeiová and Paluš, 2012). This index can be estimated as:

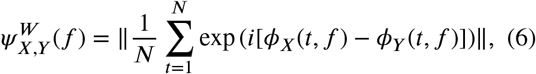

where, *ϕ*_*X*_(*t, f* ) and *ϕ*_*Y*_ (*t, f* ) denote the instantaneous phases of signals *X* and *Y* respectively at time *t* and frequency *f* .

Alternatively, Mean Phase Coherence can be directly estimated from wavelet coefficients in the following manner:

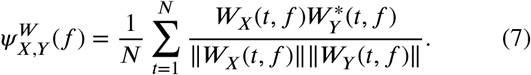

Weighted Imaginary Mean Phase Coherence or WIMPC can be obtained from the above in the same manner as WIC is estimated from Coherence:

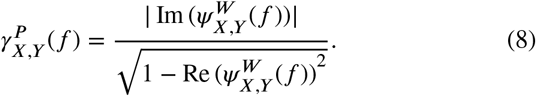

Note that a conceptually similar index called ‘lagged phase synchronization’ has been proposed in Pascual-Marqui et al. (2011). However, it is computed using normalized Fourier transforms instead of normalized Wavelet transforms. Since EEG is non-stationary and we are interested in a time-resolved version of mean phase coherence, we adopt the wavelets approach for its estimation. An alternative computation technique consists of applying a Fourier transform based bandpass filter followed by the Hilbert transform to calculate the instantaneous phase. The wavelet transform provided both filtering and phase estimation. The equivalence of these approaches has been demonstrated by Le Van Quyen et al. (2001). For the consistency of estimating MPC with coherence, we reformulated the coherence (and WIC) also into wavelet domain. It is also interesting to note that Mezeiová and Paluš (2012) demonstrated that wavelet-based MPC and FFT-based coherence provided equivalent results in sleep EEG analysis.

Using 19 channel EEG recordings for each patient, WICs were computed between all 171 distinct pairs of channels. As only every 2s of EEG signal recordings were expected to be continuous after preprocessing, ‖*W*_*X*_ (*f*) ‖, ‖*W*_*Y*_ (*f*) ‖ and*W*_*X,Y*_ (*f* ), were computed from 2s or equivalently 500 samples length signals and then summed up over consecutive 8 windows of 500 samples length to yield coherences and consequently WICs over 4000 samples or 16s length segments. Similarly, mean phase coherences were also computed from 500 samples length segments and then its average computed from 8 such consecutive windows. Finally, WIMPC for 16s segments was estimated based on these averaged estimates. To assess the significance of WICs as well as WIMPCs at each scale for each segment, 30 Fourier transform surrogates were generated for each EEG channel and corresponding WICs (WIMPCs) computed from each set of surrogates in the same manner as for the original signals. WIC (WIMPC) between original signals (for a particular scale and segment) was considered to be significant if the z-score of the original value against the population of surrogates was found to be greater than a threshold of 1.65.

Further, the significant WICs (WIMPCs) computed for 78 different scales (in the range of 1-40 Hz, with a step size of 0.5 Hz) were aggregated by summing them within predefined frequency ranges to give the WICs (WIMPCs) associated with 6 standard EEG bands. These were delta (*δ*) [1.5 - 3.5 Hz], theta (*θ*) [4 - 7.5 Hz], alpha (*α*) [8 -12Hz], beta1 (*β*_1_) [12.5 - 17.5 Hz], beta2 (*β*_2_) [18 - 25.5Hz] and gamma (*γ*) [26 - 40 Hz]. WICs and WIMPCs from 16s length segments associated with the 6 frequency bands were further used to construct functional brain networks for each frequency band separately. They were then used for distinguishing between the responders and non-responders using the features described below.

### 2.3.2. Cross-Hemispheric Connectivity

Cross-hemispheric (CH) connectivity was estimated for all the 6 frequency bands using both WICs and WIMPCs. To compute average **weighted CH connectivity** (*CH*^*W*^ ), the significant WICs (WIMPCs) estimated for all connections between the left and the right hemisphere were averaged across all the 16s segments available in the patient’s recording for a particular session. Basically, in the computation of CH connectivity, all intra-hemispheric connections and any connections of channels on the midline were ignored.

In order to compute average **binary CH connectivity** (*CH*^*B*^) across segments, all non-zero significant WICs (WIMPCs) for connections between the left and the right hemisphere were counted for all segments and divided by the number of segments available for that patient for a particular session.

#### 2.3.3. Lateral Asymmetry

Lateral asymmetry (LA) or left-right asymmetry was estimated for coherences computed for the 6 frequency bands separately as in the case of CH connectivity. Again, a weighted as well as a binary index was computed.

The difference in the strength of symmetrically placed links in the left and right hemisphere were compared. **Absolute weighted LA** was estimated as below:

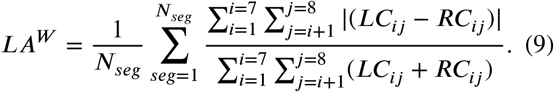

*LC*_*ij*_, *RC*_*ij*_, *seg, N*_*seg*_ are as defined in Definition 1 of LA. To understand the index estimated intuitively, we basically quantify the difference in the weights of symmetrically placed links in the left and the right hemisphere. The sum of the absolute value of this quantity is taken for all possible symmetric links. This quantity is normalized by the sum of all intra-hemispheric links considered. Finally, we obtain the average of this quantity across all time segments.

**Absolute binary LA** was estimated as follows:

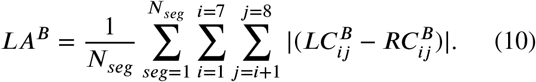

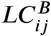 refers to the binary link between *i*^*th*^ and *j*^*th*^ electrode in the left hemisphere and 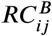 to the corresponding link in the right hemisphere. In this case, average asymmetry in the existence of symmetrically placed links in the left and right hemisphere is quantified.

## 3. Results

### 3.1. CH connectivity in response to anti-depressant treatment

Weighted CH connectivity as well as binary CH connectivity was estimated for responders and non-responders for Visit 1, i.e. before the start of the treatment (*CH*_*V is*1_) as well as for Visit 2, i.e. one week after the start of the treatment (*CH*_*V is*2_). Since there are multiple factors (response, treatment, and visit) that could lead to a change in CH connectivity and one of them is a repeated factor (the EEG recording is performed twice on the same patient), we performed statistical analysis on the data with repeated measures analysis of variance (ANOVA) (Davis et al., 2002). The repeated measures ANOVA design consists of two independent between-subject factors — response and treatment, and one dependent within-subject factor — visit, each having two levels. The response factor has the levels of responders and non-responders to treatment, while the treatment factor has the pharmacological and neurostimulation levels. The within-subject factor, visit, has two levels — visit 1 and visit 2. The model incorporates the three individual factors as well as all their interactions i.e., the three pairwise interactions and the single three-way interaction. Separate analyses are conducted for each EEG frequency band. We note that the treatment factor is not central to our analysis, and is primarily included in the model to control for any possible effects attributable to the two different therapies. In Section 4 we elaborate on how the results change when this factor is not taken into account.

Table 2a and Table 3a show the results of the above described ANOVA procedure for weighted and binary CH connectivity respectively estimated using weighted imaginary coherence. False Discovery Rate (FDR) adjusted *p*-values for the six different frequency bands are shown in the tables. These are obtained using the Benjamini–Yekutieli FDR control procedure (Benjamini and Yekutieli, 2001). Significant *p*-values (< 0.05) are in bold font.

**Table 2a.**
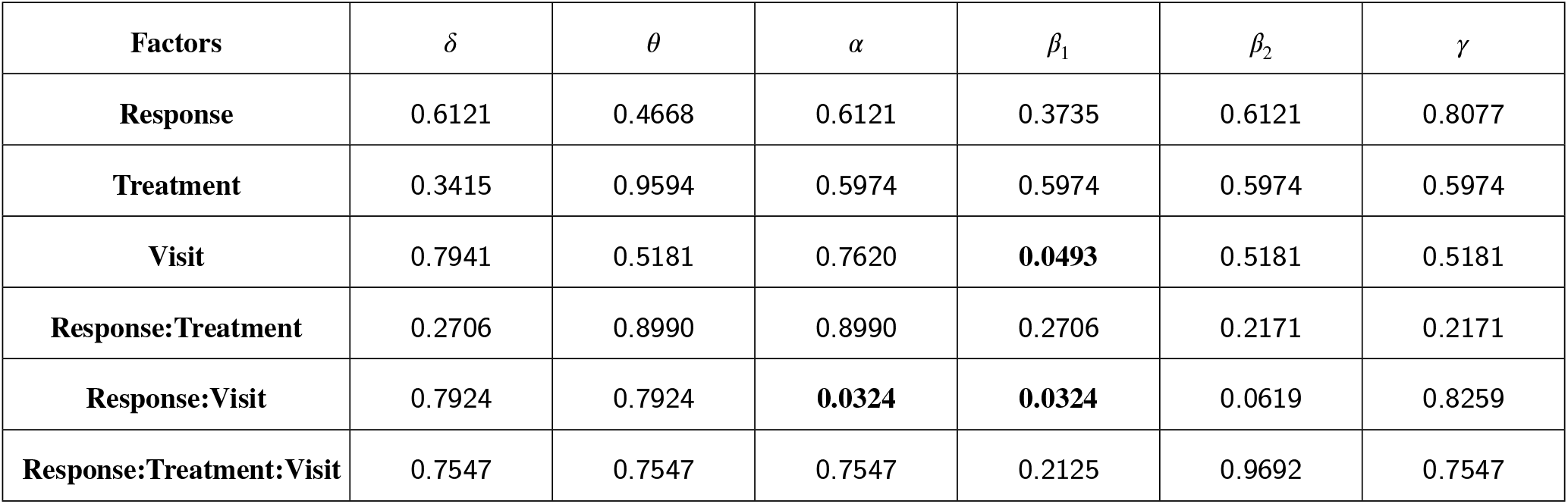
FDR adjusted *p*-values obtained from a repeated measures ANOVA test on weighted cross-hemispheric connectivity estimated using weighted imaginary coherence for six different frequency bands.

| Factors | $\delta$ | $\theta$ | $\alpha$ | $\beta_1$ | $\beta_2$ | $\gamma$ |
| --- | --- | --- | --- | --- | --- | --- |
| Response | 0.6121 | 0.4668 | 0.6121 | 0.3735 | 0.6121 | 0.8077 |
| Treatment | 0.3415 | 0.9594 | 0.5974 | 0.5974 | 0.5974 | 0.5974 |
| Visit | 0.7941 | 0.5181 | 0.7620 | **0.0493** | 0.5181 | 0.5181 |
| Response:Treatment | 0.2706 | 0.8990 | 0.8990 | 0.2706 | 0.2171 | 0.2171 |
| Response:Visit | 0.7924 | 0.7924 | **0.0324** | **0.0324** | 0.0619 | 0.8259 |
| Response:Treatment:Visit | 0.7547 | 0.7547 | 0.7547 | 0.2125 | 0.9692 | 0.7547 |

**Table 2b.** Multiple pairwise comparison tests for the significant cases in Table 2a.

| Significant case considered | Group1, Group 2 | Response type | Difference | $p$-value | Cohen's d |
| --- | --- | --- | --- | --- | --- |
| Visits, $\beta_1$ band | Visit1, Visit2 | Both | 3.1124 | 0.0075 | 0.1502 |
| Response:Visits, $\alpha$ band | Visit1, Visit2 | Responders | 6.1898 | 0.0513 | 0.1351 |
|  |  | Non-Responders | -4.2610 | 0.0911 | -0.1304 |
| Response:Visits, $\beta_1$ band | Visit1, Visit2 | Responders | **6.3564** | **0.0005** | **0.2778** |
|  |  | Non-Responders | -0.1315 | 0.9275 | 0.0426 |

**Table 3a.** FDR adjusted *p*-values obtained from a repeated measures ANOVA test on binary cross-hemispheric connectivity estimated using weighted imaginary coherence for six different frequency bands.

| Factors | $\delta$ | $\theta$ | $\alpha$ | $\beta_1$ | $\beta_2$ | $\gamma$ |
| --- | --- | --- | --- | --- | --- | --- |
| Response | 0.6784 | 0.5592 | 0.4254 | 0.1759 | 0.4821 | 0.6784 |
| Treatment | 0.0898 | 0.8111 | 0.5481 | 0.5481 | 0.5481 | 0.5930 |
| Visit | 0.7650 | 0.7650 | 0.7650 | 0.1957 | 0.7650 | 0.6527 |
| Response:Treatment | 0.1286 | 0.7516 | 0.7516 | 0.3513 | 0.1286 | 0.1286 |
| Response:Visit | 0.7293 | 0.7293 | 0.0716 | **0.0007** | 0.1012 | 0.9335 |
| Response:Treatment:Visit | 0.5929 | 0.7271 | 0.8433 | 0.5768 | 0.8433 | 0.8433 |

**Table 3b.** Multiple pairwise comparison tests for the significant cases in Table 3a.

| Significant case considered | Group1, Group 2 | Response type | Difference | *p*-value | Cohen's d |
| --- | --- | --- | --- | --- | --- |
| Response:Visits, $\beta_1$ band | Visit1, Visit2 | Responders | **2.6724** | **0.0001** | **0.3050** |
|  |  | Non-Responders | -0.7814 | 0.1522 | -0.0668 |

The ANOVA test can indicate significant factors but cannot always specify which group means have changed (e.g., when there is significant interaction terms). Further post-hoc analysis was performed for the cases where the obtained *p*-values are significant. Tukey’s Honestly Significant Difference test for performing multiple pairwise comparisons helped identify which specific groups differ significantly while controlling the family-wise error rate (FWER). Tables 2b and 3b illustrate the results for this analysis, presenting the difference of CH connectivity between groups, *p*-values as well as a Cohen’s *d* value to quantify the effect size for each of the cases deemed significant in Tables 2a and 3a respectively. For cases where *p*-value < 0.05 and Cohen’s *d* > 0.2, results are in bold font.

For the networks computed using mean phase coherence information, that is, using WIMPCs, the quantities *CH*_*V is*2_ and *CH*_*V is*1_ were once again computed for all the subjects. Repeated measures ANOVA test was performed in the same way as before and results for the weighted case are given in Table 4a and for the binary case in Table 5a. Post-hoc results are summarized in tables 4b and 5b respectively.

**Table 4a.** FDR adjusted *p*-values obtained from a repeated measures ANOVA test on weighted cross-hemispheric connectivity estimated using weighted imaginary mean phase coherence for six different frequency bands.

| Factors | $\delta$ | $\theta$ | $\alpha$ | $\beta_1$ | $\beta_2$ | $\gamma$ |
| --- | --- | --- | --- | --- | --- | --- |
| **Response** | 0.6557 | 0.3014 | 0.5687 | 0.3014 | 0.5687 | 0.6262 |
| **Treatment** | 0.2193 | 0.9741 | 0.6826 | 0.6826 | 0.6826 | 0.6826 |
| **Visit** | 0.7420 | 0.4426 | 0.4426 | 0.0724 | 0.4426 | 0.4426 |
| **Response:Treatment** | 0.2804 | 0.9050 | 0.9633 | 0.4657 | 0.2804 | 0.2804 |
| **Response:Visit** | 0.8303 | 0.5542 | **0.0361** | 0.0684 | **0.0361** | 0.8303 |
| **Response:Treatment:Visit** | 0.9126 | 0.7679 | 0.7679 | 0.3580 | 0.7679 | 0.7679 |

**Table 4b.** Multiple pairwise comparison tests for the significant cases in Table 4a.

| Significant case considered | Group1, Group 2 | Response type | Difference | *p*-value | Cohen's d |
| --- | --- | --- | --- | --- | --- |
| Response:Visits, $\alpha$ band | Visit1, Visit2 | Responders | 6.4640 | 0.0281 | 0.1484 |
|  |  | Non-Responders | -3.0768 | 0.1881 | -0.1146 |
| Response:Visits, $\beta_2$ band | Visit1, Visit2 | Responders | **2.2588** | **0.0096** | **0.2928** |
|  |  | Non-Responders | -0.6944 | 0.3159 | -0.0607 |

**Table 5a.** FDR adjusted *p*-values obtained from a repeated measures ANOVA test on binary cross-hemispheric connectivity estimated using weighted imaginary mean phase coherence for six different frequency bands.

| Factors | $\delta$ | $\theta$ | $\alpha$ | $\beta_1$ | $\beta_2$ | $\gamma$ |
| --- | --- | --- | --- | --- | --- | --- |
| **Response** | 0.6648 | 0.4078 | 0.4078 | 0.2340 | 0.4078 | 0.6397 |
| **Treatment** | 0.1134 | 0.8217 | 0.4287 | 0.4287 | 0.4287 | 0.4287 |
| **Visit** | 0.7929 | 0.7929 | 0.7700 | 0.2146 | 0.3360 | 0.5770 |
| **Response:Treatment** | 0.2536 | 0.7915 | 0.7915 | 0.5091 | 0.1589 | 0.1589 |
| **Response:Visit** | 0.9666 | 0.4224 | 0.1323 | **0.0088** | 0.1323 | 0.9723 |
| **Response:Treatment:Visit** | 0.9298 | 0.9298 | 0.9298 | 0.4016 | 0.9298 | 0.9419 |

**Table 5b.**
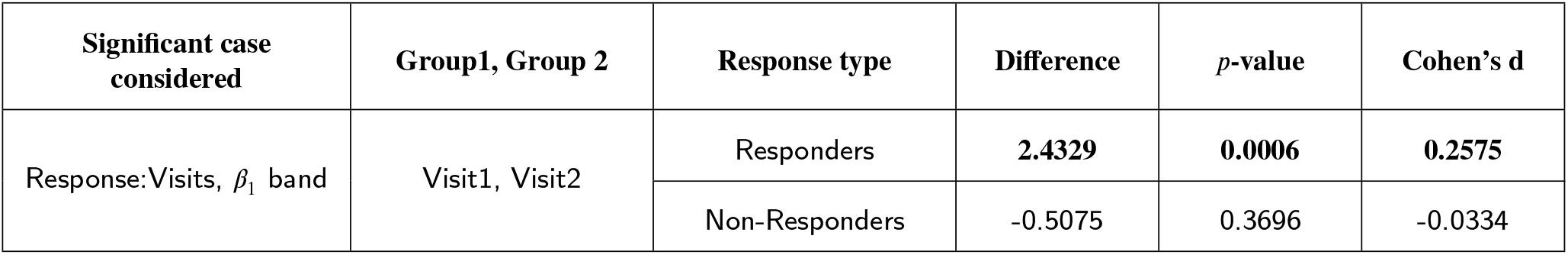
Multiple pairwise comparison tests for the significant cases in Table 5a.

| Significant case considered | Group1, Group 2 | Response type | Difference | *p*-value | Cohen's *d* |
| --- | --- | --- | --- | --- | --- |
| Response:Visits, $\beta_1$ band | Visit1, Visit2 | Responders | **2.4329** | **0.0006** | **0.2575** |
|  |  | Non-Responders | -0.5075 | 0.3696 | -0.0334 |

It can be observed from Tables 2a and 2b that a significant difference between the visits exists and can be primarily ascribed to a decrease in the *CH*^*W*^ in the *β*_1_ band for the responders at Visit 2 (*p*=0.005, *d*=0.2778). No such decrease was observed for the non-responders. *p*-values for the single factor ‘Visit’ in *β*_1_ band and for the pairwise interaction ‘Response: Visit’ in the *α* and *β*_1_ bands are significant. However, post-hoc analysis confirms that these observed differences in the *β*_1_ band were the result of a significant difference between the visits of the responders. The difference in the *α* band marginally loses statistical significance when we perform post-hoc analysis.

For the binary case using WIC, we observe a significant difference in the *β*_1_ band again for the ‘Visit: Response’ interaction. Post-hoc analysis reveals that it arose as a result of a significant change in *CH*^*B*^ for the responders (*p*=0.0001, *d* = 0.3050) while no such change was observed for the non-responders (Tables 3a, 3b). *CH*^*B*^ changes using the WIMPC measure also conformed to these results. Tables 5a and 5b reveal that a significant *p*-value in the *β*_1_ band can be attributed to a decrease in *CH*^*B*^ for the responders at Visit 2 w.r.t. Visit 1 (*p*=0.0006, *d* = 0.2575). For *CH*^*W*^ based on WIMPC, significant *p*-values were observed for the ‘Response: Visit’ interaction alone in the *α* and *β*_2_ bands (Table 4a). Based on post-hoc analysis as shown in table 4b, it can be seen that the differences were due to a decrease in *CH*^*W*^ for the responders at Visit 2. These results were stronger for the *β*_2_ band (*p* = 0.0096, *d* = 0.2928). FDR adjusted *p*-value for the *β*_1_ band was 0.0684, slightly above the significance level of 0.05.

### 3.2. Lateral asymmetry in response to anti-depressant treatment

Analogously to the CH connectivity, weighted lateral asymmetry *LA*^*W*^ as well as binary lateral asymmetry *LA*^*B*^ was estimated for all the subjects during Visit 1 as well as Visit 2 of the treatment. Once again, a repeated measures ANOVA test with details as described in the previous subsection 3.1 was performed.

FDR adjusted *p*-values for the six different frequency bands for measures *LA*^*W*^ and *LA*^*B*^ estimated using Weighted Imaginary Coherence are displayed in Tables 6a and 7a respectively. Significant *p*-values (< 0.05) are in bold font. Post-hoc analysis to investigate the meaning of observed significant *p*-values was performed in the same way as for CH connectivities before. These results are shown in table 6b for the weighted case. No significant *p*-values were obtained for the binary case.

**Table 6a.** FDR adjusted *p*-values obtained from a repeated measures ANOVA test on weighted lateral asymmetry estimated using weighted imaginary coherence for six different frequency bands.

| Factors | $\delta$ | $\theta$ | $\alpha$ | $\beta_1$ | $\beta_2$ | $\gamma$ |
| --- | --- | --- | --- | --- | --- | --- |
| Response | 0.7912 | 0.7912 | 0.2989 | 0.2989 | 0.7912 | 0.7914 |
| Treatment | 0.3218 | 0.6485 | 0.3218 | 0.6485 | 0.6485 | 0.6485 |
| Visit | 0.5290 | 0.9362 | 0.4605 | 0.2750 | 0.2750 | 0.9362 |
| Response:Treatment | 0.6115 | 0.6491 | 0.6823 | 0.6115 | 0.6491 | 0.6115 |
| Response:Visit | 0.2738 | 0.3507 | 0.0712 | **0.0143** | **0.0455** | 0.5103 |
| Response:Treatment:Visit | 0.8530 | 0.9566 | 0.9216 | 0.5526 | 0.8530 | 0.5526 |

**Table 6b.** Multiple pairwise comparison tests for the significant cases in Table 6a.

| Significant case considered | Group1, Group 2 | Response type | Difference | $p$-value | Cohen's d |
| --- | --- | --- | --- | --- | --- |
| Response:Visits, $\beta_1$ band | Visit1, Visit2 | Responders | **-0.0393** | **0.0023** | **-0.2820** |
|  |  | Non-Responders | 0.0114 | 0.2647 | 0.0715 |
| Response:Visits, $\beta_2$ band | Visit1, Visit2 | Responders | **-0.0249** | **0.0074** | **-0.2722** |
|  |  | Non-Responders | 0.0042 | 0.5679 | 0.0321 |

**Table 7a.** FDR adjusted *p*-values obtained from a repeated measures ANOVA test on binary lateral estimated using weighted imaginary coherence for six different frequency bands.

| Factors | $\delta$ | $\theta$ | $\alpha$ | $\beta_1$ | $\beta_2$ | $\gamma$ |
| --- | --- | --- | --- | --- | --- | --- |
| Response | 0.8746 | 0.8746 | 0.8746 | 0.8746 | 0.8746 | 0.8746 |
| Treatment | 0.6314 | 0.5003 | 0.1300 | 0.5174 | 0.9893 | 0.5733 |
| Visit | 0.5686 | 0.9216 | 0.9216 | 0.9216 | 0.9216 | 0.9216 |
| Response:Treatment | 0.6078 | 0.7270 | 0.7270 | 0.7270 | 0.9776 | 0.7270 |
| Response:Visit | 0.8139 | 0.8139 | 0.8139 | 0.8139 | 0.8139 | 0.8139 |
| Response:Treatment:Visit | 0.8567 | 0.8567 | 0.8567 | 0.8567 | 0.8567 | 0.8567 |

Finally, for the same analysis using the WIMPC measure, the results are in Tables 8a, 8b for the weighted case and Table 9a for the binary case.

**Table 8a.** FDR adjusted *p*-values obtained from a repeated measures ANOVA test on weighted lateral asymmetry estimated using Weighted Imaginary Mean Phase Coherence for six different frequency bands.

| Factors | $\delta$ | $\theta$ | $\alpha$ | $\beta_1$ | $\beta_2$ | $\gamma$ |
| --- | --- | --- | --- | --- | --- | --- |
| Response | 0.8918 | 0.8347 | 0.6427 | 0.3700 | 0.8918 | 0.8918 |
| Treatment | 0.2243 | 0.8104 | 0.2243 | 0.8104 | 0.8104 | 0.8104 |
| Visit | 0.3768 | 0.4702 | 0.6960 | 0.2171 | 0.1585 | 0.7901 |
| Response:Treatment | 0.7943 | 0.7943 | 0.7943 | 0.7943 | 0.7943 | 0.7943 |
| Response:Visit | 0.1534 | 0.1534 | 0.2071 | **0.0132** | 0.2071 | 0.6684 |
| Response:Treatment:Visit | 0.9818 | 0.9818 | 0.9818 | 0.1288 | 0.9818 | 0.8711 |

**Table 8b.** Multiple pairwise comparison tests for the significant cases in Table 8a.

| Significant case considered | Group1, Group 2 | Response type | Difference | $p$-value | Cohen's d |
| --- | --- | --- | --- | --- | --- |
| Response:Visits, $\beta_1$ band | Visit1, Visit2 | Responders | **-0.0393** | **0.0017** | **-0.2429** |
|  |  | Non-Responders | 0.0104 | 0.2952 | 0.0493 |

**Table 9a.**
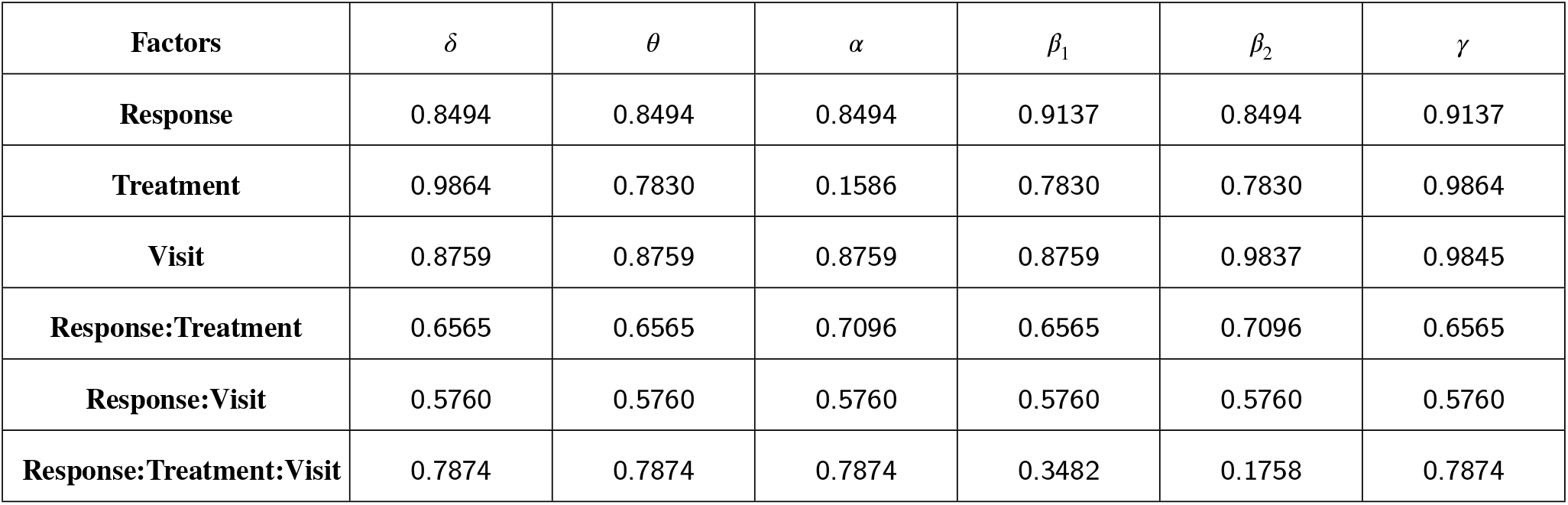
FDR adjusted *p*-values obtained from a repeated measures ANOVA test on binary lateral asymmetry estimated using weighted imaginary mean phase coherence for six different frequency bands.

No significant differences were observed in *LA*^*B*^ computed based on WIC as well as WIMPC for any of the factors as can be seen from Tables 7a and 9a. For *LA*^*W*^ computed using WIC as well as WIMPC, significant *p* values were observed for the interaction factor ‘Response:Visit’ in the *β*_1_ band and can be attributed to an increase in *LA*^*W*^ at Visit 2 w.r.t. Visit 1 for the responders (Tables 6a, 6b, 8a, 8b). Additionally, this increase was also observed in the *β*_2_ band for the estimates obtained using WIC (Tables 6a, 6b).

## 4. Discussion and Conclusions

EEG-derived measures are increasingly investigated as potential neurophysiological markers in neuroscience and psychiatry because EEG is low-cost, non-invasive, and widely available. In this study, we are interested in depression which has become a major cause of concern around the globe and wanted to track neurophysiological indicators which can act as early predictors of response to antidepressant treatment. As depression treatment often takes many weeks, waiting for the treatment course duration to observe the effect of treatment is both a waste of time and resources, and also holds a risk of further deterioration of patient’s health. Hence, we wanted to track changes in EEG based brain connectivity one week after treatment is started for the patients.

While some studies have focused on the development of such biomarkers for MDD, neither of them is close to being used in the clinic. Some of the reasons for this is because these studies are often limited to particular treatment types (Oakley et al., 2022; Bailey et al., 2019), done on a small sample of subjects (Olbrich et al., 2014; Bailey et al., 2019) or done with EEG acquired several weeks after the start of the treatment (Lee et al., 2011).

In this study we analyse a relatively large database of patients, comprising of 176 subjects being administered different types of treatment and focus on features based on brain connectivity networks estimated using coherence obtained from different frequency bands. Since weighted imaginary coherences and mean phase coherences are used, which are known to be resistant to spurious connectivities arising from EEG volume conduction effects, we expect the network estimation to be reliable, capturing only the ‘true’ connectivities (Paluš et al., 2023).

In the comprehensive results that we present, the two features analysed were lateral asymmetry and cross-hemispheric connectivity in 6 different frequency bands. Further, these features were estimated from both weighted and binary brain networks. In the latter case, the networks were binarised based on significance testing with respect to surrogate distributions. A repeated measures ANOVA test was performed to check for differences in brain connectivity (based on the measures considered) at Visit 1 and Visit 2 for all the subjects. Since multiple factors could lead to a change in connectivity, the test design consists of two independent between-subject factors — response and treatment type, and one dependent within-subject factor — visit. We found that the interaction of ‘response’and ‘visit’leads to a significant change in connectivity in certain frequency bands for the measures considered. The individual factors or other interaction terms were not found to have substantially significant *p*-values. Post-hoc analyses tests for multiple pairwise comparisons revealed that these observed significant *p*-values were a result of a change in connectivity of the responders at Visit 2 w.r.t. Visit 1. Interestingly, non-responders did not exhibit any significant change in connectivity for the measures considered.

The treatment factor did not yield statistically significant main or interaction effects for our dataset. Hence, there was no statistically significant evidence that the main findings differed by treatment modality. This is itself interesting since the two therapies involve quite different mechanisms. As expected, removal of this factor from the ANOVA model did not substantially change the results and the derived conclusions.

Hence, the above analysis reveals a couple of promising indicators of neurophysiological response. Based on CH connectivity: *CH*^*W*^ based on WIC in the *β*_1_ band; *CH*^*B*^ based on WIC and WIMPC in the *β*_1_ band; *CH*^*W*^ based on WIMPC in the *α* and *β*_2_ bands. For the latter case, *CH*^*W*^ in *α* band can be said to be a weak indicator as the Cohen’s *d* for the pairwise comparison in this case < 0.2 (Table 4b). For *CH*^*W*^ based on WIMPC in the *β*_1_ band, the FDR corrected *p*-value for the Response:Visit interaction factor is found to be not significant but is quite close to 0.05 (*p* = 0.0684). One reason why in this case significance is lost is because for weighted connectivity estimates, variance is increased and consequently sensitivity decreased. Based on lateral asymmetry: *LA*^*W*^ in the *β*_1_ band based on both WIC and WIMPC; *LA*^*W*^ in the *β*_2_ band based on WIC. For binary lateral asymmetry, no significant differences in any frequency band were observed as for most of the subjects this measure was often estimated to be zero. It is usually the case that symmetric links are present (or not present) in both hemispheres at the same time. Although the weights of connections (in the left and right hemispheres) may be a symmetric and of importance, the binarised networks were often symmetric and hence did not provide much information.

Overall, *β*_1_ band seemed to be of consistent importance when observing changes in brain connectivity of responders at 1 week post start of treatment - both CH and LA differences were observed in this band. Additionally, binary CH connectivity and and weighted lateral asymmetry were consistently good indicators with respect to use of different brain network estimation approaches – WIC and WIMPC, meaning that there were changes observed in these measures irrespective of the use of amplitude-cum-phase or phase based approaches for construction of brain networks. Based on these observations, a decrease in binary CH connectivity and an increase in weighted lateral asymmetry in the *β*_1_ are some changes associated with an early treatment effect in responders.

There is growing evidence that age Al Zoubi et al. (2018) and sex Bučková et al. (2020) are relevant factors for EEG features. It is therefore essential to check the results after including these variables in the model. Hence, for the cases found significant with repeated measures ANOVA test above, a linear mixed-effects model was also tested with age and sex as additional factors. However, this model did not provide any significant *p*-value for these two factors. Since these factors did not influence the results, we decided to use the ANOVA test as it provides for a more succinct model based analysis.

A subset of the patient dataset considered here has already been analysed in a few studies where *cordance* and *alpha asymmetry* were explored as early predictors of treatment response (Bares et al., 2010, 2015b,a, 2019). In these studies, it was found that the responders show a decrease in prefrontal theta cordance, while the non-responders show an increase in occipital alpha asymmetry (Bares et al., 2019). Since these power spectral features have already been studied previously, we focused on exploring functional brain connectivity based features as predictors. Neurobiological significance of EEG connectivity changes in early antidepressant treatment is supported by previous research (Bareš et al., 2007, 2008; Bares et al., 2010, 2015b,a; Bareš et al., 2017; Bares et al., 2019).

Some existing studies looking at functional and structural connectivity changes during depression have used lateral asymmetry and cross-hemispheric connectivity like formulations as features and observed some modifications of these in depression patients (Quraan et al., 2014; Hermesdorf et al., 2016). The argument behind using these as features is to observe the hampering of neural oscillations synchrony during disorders which are crucial in maintaining information integration in the healthy brain. Thus we focused on these features as potential early indicators of treatment response.

In our study, a decrease in binary cross-hemispheric connectivity and an increase in weighted lateral a symmetry in the *β*_1_ band were associated with early response to treatment. These findings should be regarded as candidate neurophysiological correlates of early treatment response rather than clinically established biomarkers, and they require validation in independent cohorts. These changes observed in the responders to anti-depressant treatment are robust across the connectivity measures examined, as they were not found to be influenced by the particular type of treatment administered (pharmacological or neurostimulation) or by the measure used to construct brain networks (phase or amplitude-cum-phase based). They are potentially a result of changes observed in cortical excitability and network reorganization associated with treatment response. We would like to note that interpretation of our findings should remain cautious because response status was defined after 4 weeks whereas EEG change was measured after 1 week, treatment modalities were heterogeneous, and the present findings have not yet been externally validated.

Future work would entail testing on an independent set of patients and also observing if the change in CH/ LA connectivity arose as a result of changes in some specific connections. Preliminary analysis for patients showing the highest reduction in CH connectivity showed that connections with the temporal cortex (of frontal, parietal and occipital regions from the opposite hemisphere) were reduced. Also, preliminary analysis of patients with an increase in LA showed that connections with the temporal regions contributed the most to observed left-right asymmetry. However, this finding was not consistent across all responders. Extensive analysis will be done to check what is the predominant cause of decreased binary CH connectivity in the *β*_1_ band and increased weighted LA in the *β*_1_ band. We also plan to study changes in CH and LA connectivity in functional networks constructed using directed causality measures and see if such analysis can help to give any insights on the reasons for change in CH/ LA connectivity observed in this work. Another important task for future work is to test for the predictive accuracy of the neurophysiological indicators observed in this study.

## Acknowledgements and Funding Sources

This work was supported by the Czech Science Foundation (Project No. GF21-14727K and No. 23-07074S), the Czech Academy of Sciences through the Praemium Academiae award to M. Paluš, the European Regional Development Fund Project Brain Dynamics (Project No. CZ.02.01.01/00/22_008/0004643), and the Charles University Cooperatio Neurosciences Research Program.

## CRediT authorship contribution statement

**Aditi Kathpalia**: Methodology, Software, Validation, Formal Analysis, Investigation, Data Curation, Writing - original Draft, review and editing, Visualization. **Ioannis Vlachos**: Methodology, Software, Validation, Writing - review and editing. **Jaroslav Hlinka**: Methodology, Supervision, Writing - review and editing. **Martin Brunovský**: Resources, Data Curation, Writing - review and editing, Funding acquisition. **Martin Bareš**: Resources, Data Curation. **Milan Paluš**: Conceptualization, Methodology, Software, Validation, Writing - review and editing, Supervision, Project administration, Funding acquisition.

## Footnotes

1 The terminology of ‘true’ physiological connectivity has been used in Nolte et al. (2004) and subsequent articles. Here, it does not refer to mechanistic estimation of connectivity but functional connectivity itself which is devoid of any confounding effects, such as those arising from volume conduction in EEG.

